# TCF4 and CtBP1 Repress *LEF1* to Maintain a TCF4-centric Transcriptional Program in Colon Cancer

**DOI:** 10.64898/2026.09.25.751839

**Authors:** Markus A. Brown, Sanam Yaghoubi, Juliane Kamlah, Wei-Dong Chen, Darawalee Zong, Sepehr Talebian Khorasani, Tianzuo Zhan, Michael Boutros, Jordi Camps, Paul Meltzer

## Abstract

The canonical WNT signaling pathway guides cell growth throughout the human lifespan. In adults, WNT plays an essential role in tissue homeostasis by maintaining tissue stem cells. In the intestine, the WNT transcription factor, TCF4, is necessary for stem cell maintenance. Aberrant activation of WNT signaling is generally accepted as an initiating event in colon cancer, leading to constitutive TCF4 activity. To investigate the role of this key transcription factor in global gene regulation, we abrogated TCF4 expression in colon cancer cells. Loss of TCF4 function resulted in the over-expression of the WNT transcription factor, *LEF1*. LEF1 was transcriptionally competent and over-compensated for TCF4 (*TCF7L2*) silencing in a WNT reporter assay. TCF4 enlisted the transcriptional repressor CtBP1, and bound the *LEF1* promoter, indicative of direct repression. TCF4 and LEF1 drove different transcriptional programs, with TCF4 favoring MYC and cell cycle progression, while LEF1 favored immune-related gene signatures. TCF4 was the primary regulator of *MYC* expression, yet also the mediator of *LEF1* repression, suggestive of competition centered on TCF4 transcriptional output despite the β-catenin saturated colon cancer nucleus. We conclude that TCF4 has dual activator-repressor activity in colon cancer.

## INTRODUCTION

Humans possess four canonical WNT signaling transcription factors: TCF1, TCF3, TCF4, and LEF1, encoded by the *TCF7*, *TCF7L1*, *TCF7L2*, and *LEF1* genes, respectively (*1–6*). The expression pattern of these factors varies by developmental stage and cell type, reflecting the crucial roles they fulfill in development and tissue homeostasis (*7*, *8*). Knockout-based functional studies have delineated largely distinct roles for the TCF/LEF family members: TCF1 mediates thymocyte differentiation (*9*), TCF3 is required for proper anterior-posterior axis induction and maintains skin stem cells (*10*, *11*), TCF4 maintains colonic epithelial stem cells (*3*, *12*), and LEF1 controls hair and tooth development (*13*). However, TCF1 and LEF1 have been shown to play partially redundant roles in lymphoid development (*14–16*), while TCF4 supports the role of TCF3 in maintaining skin stem cells (*17*), demonstrating that WNT factor co-expression can provide a layer of redundancy or an additional, supporting function.

The activity of the canonical WNT transcription factors is determined by the properties of their binding partner(s) as the TCF/LEF factors do not contain a transcriptionally competent domain. The TCF/LEF factors have binding sites for β-catenin (*18*), a potent transcriptional activator, as well as for the TLE family of transcriptional repressors (*19*, *20*). Sequence-specific DNA binding is conferred by a highly conserved HMG (high mobility group) box domain (*1*, *2*). In the absence of WNT signaling, TLE binds to the TCF/LEF factors and represses WNT target gene expression. In contrast, active WNT signaling results in high levels of nuclear β-catenin and a switch to WNT target gene expression (*21*). WNT signaling output is therefore contingent upon the binding partners present in the nucleus.

In colon cancer, WNT signaling is constitutively active. The most frequent cause is truncation of APC, whose function is to facilitate the degradation of β-catenin in the cytoplasm (*22*, *23*). Hampered degradation results in accumulation of cytoplasmic β-catenin, which migrates to the nucleus and binds TCF4, leading to constitutive expression of WNT target genes such as *MYC* (*24–27*). Sustained activity of the WNT target genes contributes substantially to neoplasia in the colon (*28–30*).

The TCF/LEF expression pattern differs between normal colon and colon cancer (*31*). In the normal colon, *TCF7L2* is highly expressed, while the expression of *TCF7* and *LEF1* is low. In colon cancer, *TCF7* and *LEF1* are expressed to a greater degree, while *TCF7L2* remains the dominantly expressed transcription factor. How WNT transcription factors balance their effects on the colon cancer transcriptional profile has remained poorly explored, especially regarding *LEF1*. This study investigated the roles of TCF4 and LEF1 on colon cancer transcriptional dynamics as well as the mechanism of repression of *LEF1* by TCF4.

## RESULTS

### *TCF7* and *LEF1* Expression is Elevated in Colon Cancer

To better understand the expression pattern of the WNT signaling transcription factors in the non-diseased human colon, RNA sequencing data in transcripts per million (TPM) from GTEx was used. Data which passed filtering metrics (see Methods) were selected. The expression ratio of the WNT signaling transcription factors in a representative subset of non-diseased colon samples was plotted (Fig. 1A). *TCF7L2*, the gene encoding TCF4, is highly and broadly expressed. *TCF7L1* is also expressed to a high degree in ∼85% of the samples. *TCF7* and *LEF1*, are either expressed to a low degree or not expressed. To determine the degree and scope of WNT transcription factor expression in colon cancer, RNA sequencing data (TPM) from TCGA was retrieved. Data which passed our filtering metrics were selected, and a representative subset was plotted (Fig. 1B). *TCF7L2* was again the dominant WNT transcription factor, while *TCF7L1* expression was only detected in 2% of the samples in contrast to its broad expression in non-diseased colon tissue. The expression of *TCF7* and *LEF1* increased relative to the expression of *TCF7L2* and are expressed in 85% and 35% of colon cancer samples, respectively. Immune infiltrates were not a confounding factor in the analysis regarding the expression of *TCF7* or *LEF1*, given their role in lymphoid development (Fig. S1). Taken together, we find that *TCF7L2* is the dominant WNT transcription factor in non-diseased colon and colon cancer. *TCF7* and *LEF1* are not expressed appreciably throughout non-diseased tissue, but are expressed in colon cancer, while *TCF7L1* is expressed in the non-diseased colon but only rarely in colon cancer.

**Figure 1.**
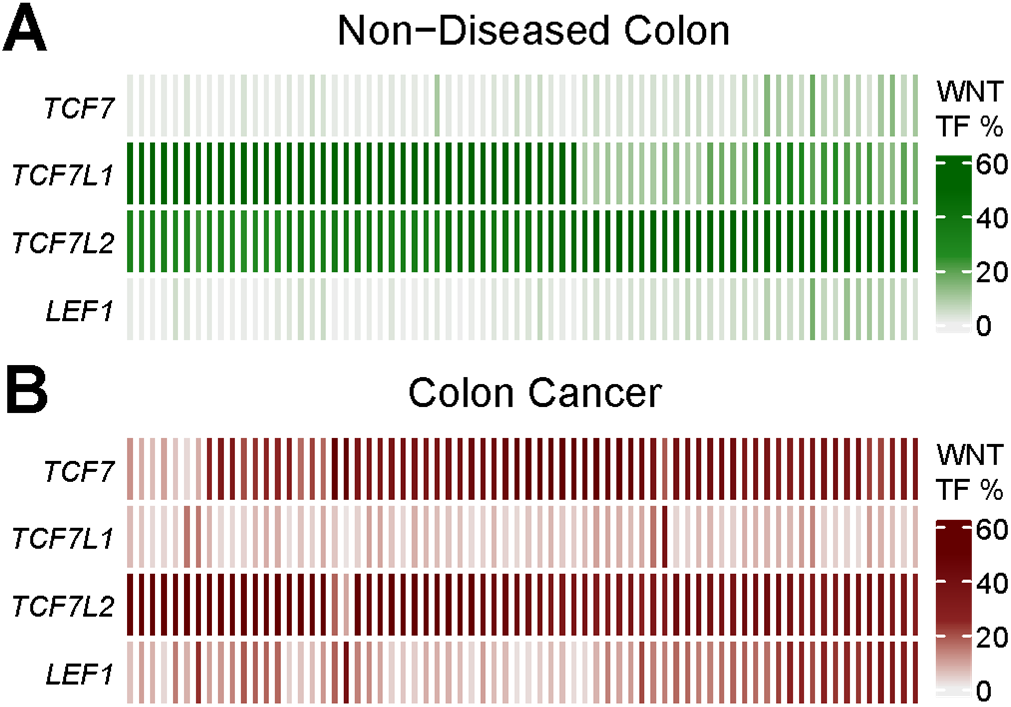
*TCF7* and *LEF1* Expression is Elevated in Colon Cancer. **| A |** Non-diseased colon RNA sequencing data in transcripts per million (TPM) from GTEx was used to generate the WNT transcription factor percentage, which is the contribution of each WNT transcription factor to the total WNT transcription factor expression per sample. The WNT transcription factor percentages in non-diseased colon tissue were plotted with the four WNT transcription factors as rows and samples as columns (n=70). **| B |** The WNT transcription factor percentage from TCGA colon cancer RNA sequencing data (TPM) was plotted with the four WNT transcription factors as rows and samples as columns (n=70).

### *TCF7L2* Silencing Results in *LEF1* Up-Regulation

Since *TCF7L2* is the dominant WNT transcription factor in colon cancer, the influence of *TCF7L2* on the global gene expression profile in the SW480 colon cancer cell line was determined with an RNAi silencing time course utilizing an siRNA targeting the 3’ end of *TCF7L2* transcripts. The siRNA decreased *TCF7L2* transcript abundance 4-fold and reduced nuclear TCF4 protein abundance 97% by the last time point (Fig. 2A, B). RNA sequencing over the time course demonstrated a progressive reduction in *TCF7L2* transcript abundance, while *LEF1* expression increased nearly 3-fold by the last time point (Fig. 2C). *TCF7* and *TCF7L1* expression did not demonstrate a consistent trend in response to *TCF7L2* silencing. Nuclear LEF1 protein levels were assessed using Western blot, which also increased nearly 3-fold by the last time point (Fig. 2D). *LEF1* up-regulation was not a result of an siRNA off-target effect as three different siRNAs, each binding a different region of the *TCF7L2* transcript, yielded dose-dependent *LEF1* up-regulation (Fig. S2A). To determine whether LEF1 was transcriptionally competent, a WNT dual-luciferase reporter assay in SW480 was performed. Silencing of *TCF7L2* alone, which induces *LEF1* over-expression, resulted in an approximate 2.5-fold increase in WNT reporter activity (green), similar to the increase in LEF1 (Fig. 2E). To determine whether LEF1 was responsible for the increase in WNT reporter activity, *TCF7L2* and *LEF1* were simultaneously silenced, thereby initiating the *TCF7L2* response, while removing the *LEF1* up-regulation component, shown in purple (Fig. 2E). The increase in WNT reporter activity was nullified, demonstrating that LEF1 was responsible for the increase in reporter activity and is therefore transcriptionally competent.

**Figure 2.**
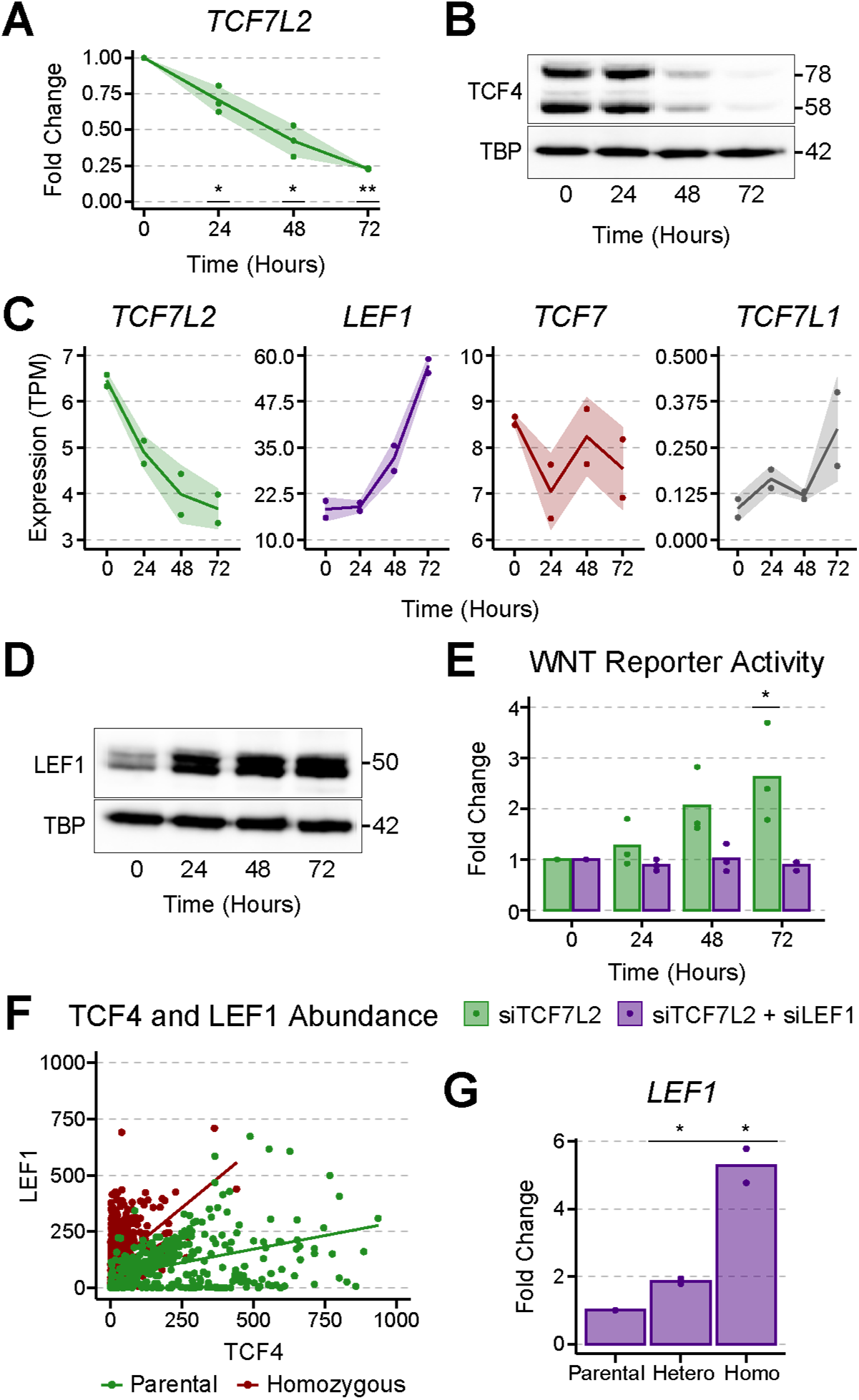
*TCF7L2* Silencing Results in *LEF1* Up-Regulation. **| A |** Quantitative PCR (qPCR) was used to assess *TCF7L2* transcript abundance upon transfection with siTCF7L2 over the time series in SW480. Three biological replicates are plotted and the ribbon denotes the standard deviation (Student’s *t*-test, * *p* < 0.05, ** *p* < 0.01). **| B |** Nuclear TCF4 abundance during the siTCF7L2 time series was determined using Western Blot with TATA-Binding Protein (TBP) as loading control. Selected molecular weights (kDa) are shown to the right. The abundance of two TCF4 isoforms decreased by ∼97% by the 72-hour timepoint. **| C |** Expression of the four WNT transcription factors upon *TCF7L2* silencing in SW480, assayed using RNA sequencing. Each dot represents a biological replicate (n=2), the line represents the average, and the shaded ribbon denotes the standard deviation. **| D |** Western blot on the nuclear protein fraction demonstrated an ∼3-fold increase in LEF1 abundance upon *TCF7L2* silencing by the 72-hour timepoint. **| E |** A dual-luciferase assay was used to determine WNT transcriptional output upon *TCF7L2* silencing, shown in green, or simultaneous silencing of *TCF7L2* and *LEF1*, shown in purple. Three biological replicates are plotted (Student’s *t*-test, * *p* < 0.05). **| F |** Ratio of TCF4 and LEF1, assayed via immunocytochemistry, from either Parental (green) or Homozygous *TCF7L2* knockout (red) lines. **| G |** *LEF1* transcript abundance was assessed using qPCR in Parental, Heterozygous, or Homozygous *TCF7L2* knockout lines. Two biological replicates are plotted (Student’s *t*-test, * *p* < 0.05).

To confirm that the TCF4-*LEF1* response occurs in a stable setting, CRISPR/Cas9 was used to remove the HMG domain of *TCF7L2* in SCDCL, a cell line generated from a treatment-naïve rectal cancer biopsy (*32*). Successful removal of the HMG domain was confirmed using PCR-based genotyping (Fig. S2B, C). *TCF7L2*-KO cells displayed marked changes in morphology and impaired growth compared to parental cells (Fig. S2D). Immunocytochemistry demonstrated an increase in the LEF1/TCF4 ratio in the homozygous *TCF7L2*-KO cells as compared to parental SCDCL (Fig. 2F). *LEF1* transcript abundance increased in heterozygous and homozygous *TCF7L2* KO clones as compared to the parental control, confirming that *LEF1* is up-regulated upon CRISPR knockout of TCF4 (Fig. 2G). Up-regulation of *LEF1* is therefore sustained upon removal of TCF4.

These data suggest that *TCF7L2* silencing results in a dose-dependent, 3-fold increase in *LEF1* expression, which translates to an equivalent increase in transcriptionally competent LEF1. The increase in LEF1 abundance over-compensates for the anticipated decrease in WNT signaling activity upon reduction of TCF4, resulting in an approximate 2.5-fold increase in WNT activity.

### Truncating APC Does Not Alter *LEF1* Expression

*TCF7L2* was then silenced in several colon cancer cell lines with varying WNT signaling characteristics to investigate whether *LEF1* up-regulation is dependent upon *APC* mutational status. The *APC* wild-type cell lines, HCT116 and LS174T, as well as the *APC* mutated lines HT29, LoVo, SW480, and DLD1 were selected. Cell lines are ordered based on the location of the earliest truncating *APC* mutation (*33*, *34*) (Fig. S3A). Neither the *APC* wild-type cell lines, HCT116 and LS174T, nor the *APC* mutated cell line, HT29 demonstrated an increase in *LEF1* expression (Fig. 3A). While HT29 harbors biallelic *APC* mutations, the second mutation occurs after the Catenin Inhibitory Domain (CID), and one allele may therefore remain functionally similar to the wild-type *APC* found in HCT116 or LS174T (*35*). The remaining *APC* mutated cell lines LoVo, SW480, and DLD1, which lack the CID, demonstrated up-regulation of *LEF1* ranging from 1.5-to 3-fold, that increased with the length of the APC fragment (Fig. 3A).

**Figure 3.**
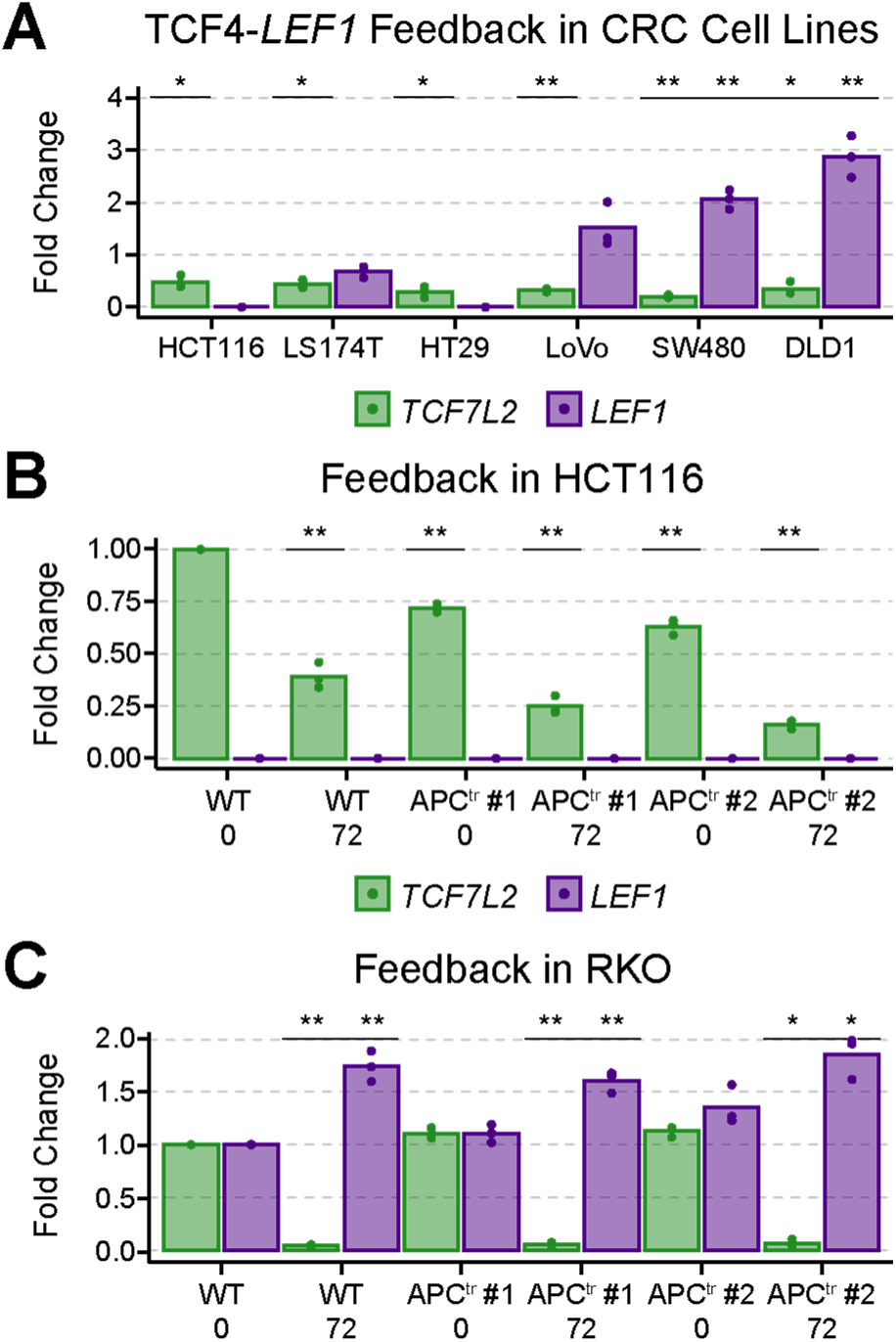
APC Does Not Alter *LEF1* Expression Patterns. **| A |** *LEF1* expression was assessed at the 72-hour timepoint using qPCR after transfection of various colon cancer cell lines with siTCF7L2. Fold change was calculated using the siNeg control for the respective cell line, set to a fold change of one (data not shown). Three biological replicates are plotted (Student’s *t*-test, * *p* < 0.05, ** *p* < 0.01). **| B |** HCT116 and two biallelically *APC*-mutated clones were transfected with siTCF7L2 for 72 hours to assess *LEF1* over-expression. Fold change was calculated using the wild-type HCT116 siNeg control (WT 0) for all samples. Three biological replicates are plotted (Student’s *t*-test, ** *p* < 0.01). **| C |** RKO and two, biallelically *APC*-mutated clones were transfected with siTCF7L2 for 72 hours to assess *LEF1* over-expression. Fold change was calculated using the wild-type RKO siNeg control (WT 0). Three biological replicates are plotted (Student’s *t*-test, * *p* < 0.05, ** *p* < 0.01).

Given the differences in *LEF1* over-expression between the various *APC* wild-type and mutated cell lines, we used two colon cancer cell lines with wild-type *APC*, HCT116 and RKO, and generated clones with biallelic, truncating mutations at the *APC* CID using CRISPR/Cas9 (*36*). The expression of *AXIN2*, *TCF7L2*, and *LEF1* was measured in the wild-type and APC-truncated lines using qPCR to determine whether *LEF1* expression was altered between the wild-type HCT116 or RKO lines and the respective *APC*-mutated counterparts. Expression of *AXIN2* increased in the truncated APC lines, demonstrating the expected increase in WNT signaling activity upon loss of APC function (Fig. S3B, D). Silencing of *TCF7L2* in APC-truncated HCT116 resulted in a 6-fold decrease in *AXIN2* expression. *LEF1* is not expressed in HCT116 and neither truncation of APC nor silencing of *TCF7L2* induced *LEF1* expression (Fig. 3B, S3C). In RKO, APC truncation resulted in an increase of *AXIN2* levels, however silencing of *TCF7L2* further increased *AXIN2* levels, demonstrating the opposite response observed in HCT116 (Fig. S3D). Importantly, silencing of *TCF7L2* in RKO led to an up-regulation of *LEF1*, which occurred largely independently of APC truncation (Fig. 3C). RKO parental cells express *LEF1* to a greater degree than the other WNT TFs (Fig. S3E). Collectively, these results demonstrate that *APC* truncation may modulate *LEF1* expression rather than altering baseline *LEF1* expression. Notably, *TCF7L2* silencing had a larger influence on *LEF1* expression than APC truncation.

### CtBP1 is the Primary Repressor of *LEF1*

To further investigate how *LEF1* expression is regulated, the role of the transcriptional repressors CtBP1 and CtBP2, which have been reported to interact with TCF4, were assessed. Hence, *CtBP1* and *CtBP2* were silenced using RNAi, and Western blot was used to assess nuclear CtBP1, CtBP2, TCF4, and LEF1 levels. Transfection with siCtBP1 or siCtBP2 led to a clear reduction in nuclear CtBP1 or CtBP2 abundance (Fig. 4A). Reduction of CtBP1 or CtBP2 did not influence TCF4 levels, removing a possible confounding factor. Intriguingly, reduction of CtBP1, but not CtBP2, resulted in a doubling of LEF1 abundance, suggesting that CtBP1 is a repressor of *LEF1*.

**Figure 4.**
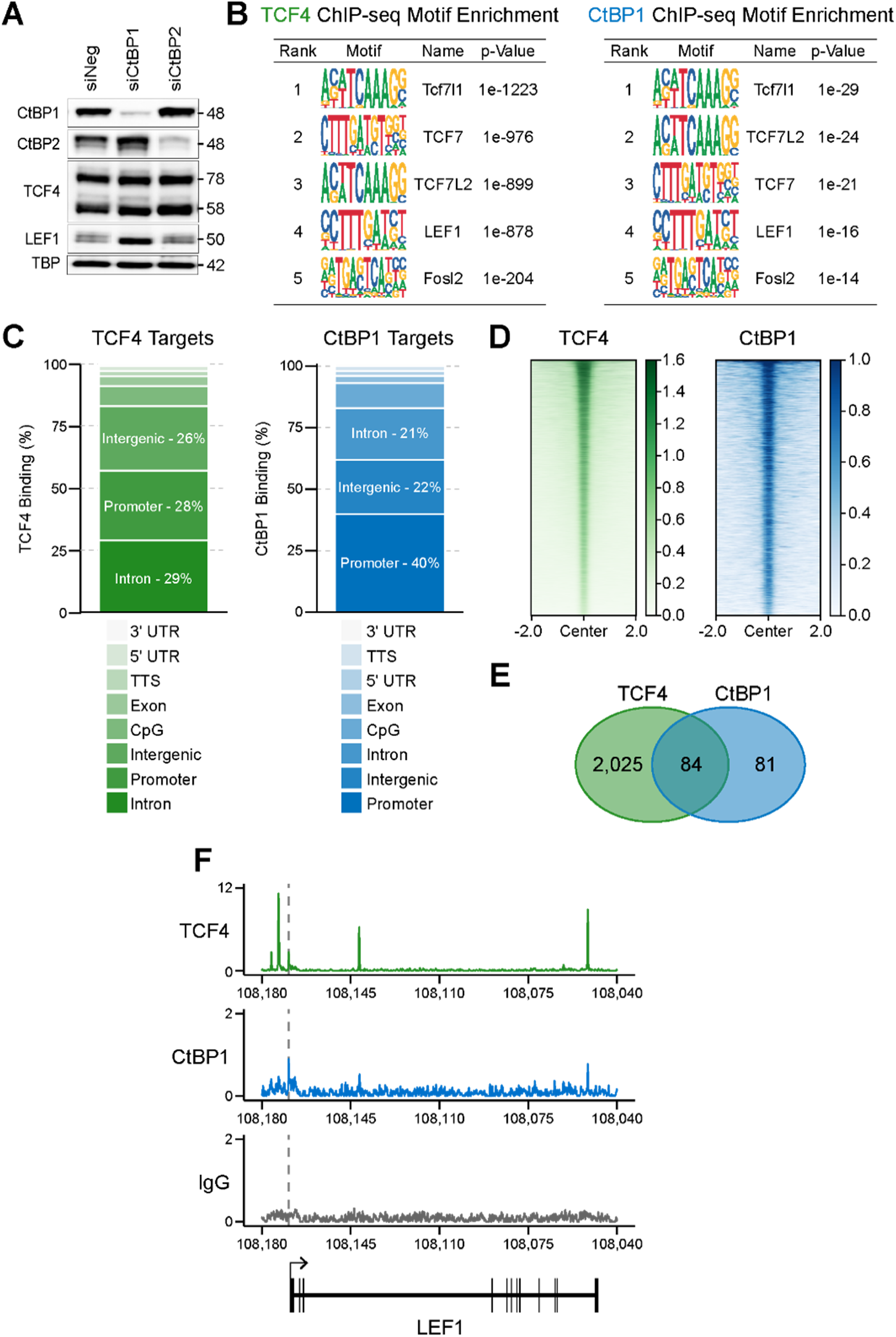
CtBP1 is the Primary Repressor of *LEF1*. **| A |** Western blot for CtBP1, CtBP2, TCF4, and LEF1 upon treatment with siNEG, siCtBP1, or siCtBP2 with TBP as loading control. Selected molecular weights (kDa) are shown to the right. **| B |** Motif enrichment from TCF4 or CtBP1 ChIP-seq using Homer. **| C |** TCF4 and CtBP1 peak distribution among genomic elements. The lists below the graphs provide the genomic elements ordered from least to most frequent. **| D |** TCF4 and CtBP1 peak heatmaps spanning 2 kilobases (kb) up- and downstream of the peak center. **| E |** Venn diagram of promoter-specific target genes from TCF4 and CtBP1. **| F |** TCF4 and CtBP1 binding at the *LEF1* locus plotted as reads per million (rpm) with genomic position in kilobases. A vertical, dashed line is plotted at the *LEF1* transcription start site (TSS). *LEF1* is located on the minus (−) strand.

To determine whether TCF4 and CtBP1 directly repress *LEF1*, we performed ChIP-Seq for TCF4 and CtBP1 in SW480 cells. The top five results from Homer motif enrichment analysis for TCF4 ChIP-seq were the four WNT transcription factors as well as Fosl2 (Fig. 4B). Interestingly, the top five motifs for CtBP1 ChIP-seq also included the four WNT transcription factors and Fosl2. TCF4 peaks were located primarily in introns or promoters, while CtBP1 peaks were located primarily in promoters, supporting the role of CtBP1 in transcriptional regulation (Fig. 4C). TCF4 peaks were predominantly narrow and sharp, consistent with its role as a sequence-specific transcription factor. CtBP1 peaks were relatively sharp, supporting the idea that CtBP1 is recruited through interactions with transcription factors, in line with its role as a transcriptional co-regulator (Fig. 4D). We identified ∼2,100 promoter-specific target genes for TCF4 and ∼160 promoter-specific target genes for CtBP1 (Fig. 4E). A large percentage of the CtBP1 target genes (∼51%) were also bound by TCF4, suggesting that TCF4 is a major channel through which the repressive effects of CtBP1 are relayed in colon cancer. TCF4 and CtBP1 exhibit overlapping peaks in the *LEF1* promoter, within a *LEF1* intron, and just preceding the 3’ UTR (Fig. 4F). Together with the RNA and protein data, these findings support a direct and cooperative mechanism of repression of *LEF1* by TCF4 and CtBP1.

### CtBP1 Represses *MYC* Expression

Given that CtBP1 appears to act as a repressor through TCF4 and that *LEF1* up-regulation is observed in multiple colon cancer cell lines, we determined the influence of CtBP1 and LEF1 on the gene expression pattern of SW480. *CtBP1* or *LEF1* expression was silenced in SW480 for 72 hours, which resulted in a 90% or 98% decrease in CtBP1 or LEF1 nuclear protein abundance, respectively (Fig. S4A, B). Bulk RNA was extracted and sequenced from *CtBP1*-silenced or *LEF1*-silenced SW480 cells in an analogous manner to *TCF7L2* silencing as performed previously. Upon silencing of *TCF7L2*, a disproportionate increase in gene expression was observed, with approximately twice the number of genes significantly up-regulated as down-regulated (Fig. 5A). Upon *LEF1* silencing the reverse trend was observed, with more genes significantly down-regulated than up-regulated. Silencing of *CtBP1* resulted in an even response with a similar number of genes significantly up- and down-regulated. This broadly suggests that the net effect of TCF4 is repressive, while the net effect of LEF1 is activating.

**Figure 5.**
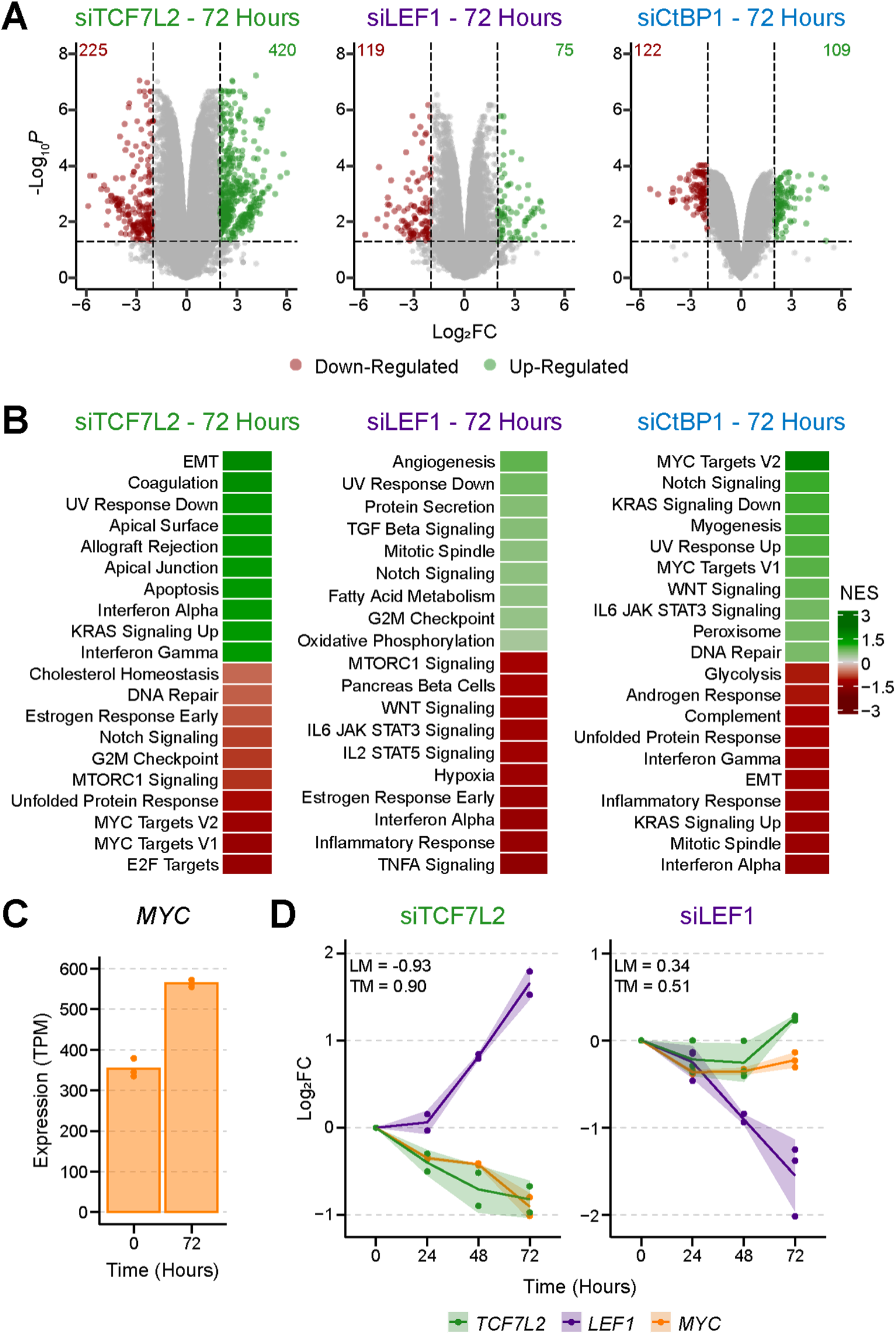
TCF4, not LEF1, Determines *MYC* Expression. **| A |** Volcano plots demonstrating the transcriptional response to silencing of *TCF7L2*, *LEF1*, or *CtBP1* in SW480 for 72 hours. The top-left and top-right numbers indicate the number of genes within the thresholds. **| B |** Gene Set Enrichment Analysis (GSEA) upon silencing of *TCF7L2*, *LEF1*, or *CtBP1* in SW480 for 72 hours from the Hallmark gene sets with the highest normalized enrichment scores (NES). **| C |** *MYC* expression upon *CtBP1* silencing for 0 (siNeg) or 72 hours (siCtBP1), shown in TPM. Three biological replicates are plotted (n=3). **| D |** Fold change in expression of *TCF7L2*, *LEF1*, and *MYC* over the siTCF7L2 or siLEF1 time series. Each dot represents a biological replicate, the line represents the average, and the shaded ribbon denotes the standard deviation. The Pearson correlation coefficients between *LEF1* and *MYC* (LM) or *TCF7L2* and *MYC* (TM) are shown within the plot areas.

To gain insight into which pathways are differentially regulated by TCF4, LEF1, and CtBP1, gene set enrichment analysis (GSEA) was performed using the hallmark gene sets from MsigDB (*37*, *38*). The top ten up-regulated and down-regulated pathways upon silencing of *TCF7L2*, *LEF1*, or *CtBP1*, based on the NES, were plotted (Fig. 5B). Silencing of *TCF7L2* resulted in the up-regulation of EMT and Immune-related pathways, while the E2F Targets and MYC Targets were down-regulated, in line with TCF4’s role as a driver of *MYC* expression and cell cycle progression. *LEF1* silencing resulted in the down-regulation of immune system gene sets, such as TNF-α Signaling, Inflammatory Response, Interferon Alpha, IL2 Signaling, and IL6 Signaling, in line with the role of LEF1 in immune system regulation (Fig. 5B). No pathways were significantly up-regulated upon *LEF1* silencing. The over-expression of *LEF1* upon *TCF7L2* silencing may explain the up-regulation of immune associated pathways, such as Interferon Alpha, in the TCF4 GSEA results. Indeed, the down-regulated pathways upon *LEF1* silencing were up-regulated upon *TCF7L2* silencing (Fig. S4C). Taken together, these results demonstrate that LEF1 can alter the expression profile of SW480 by favoring immune associated pathways.

GSEA after *CtBP1* silencing identified several significantly enriched gene sets including MYC Targets V2, KRAS Signaling Up, Interferons (Alpha and Gamma), and Epithelial-Mesenchymal Transition (Fig. 5B). Up-regulation of the MYC target gene sets suggests that CtBP1 represses *MYC*, a notable target of TCF4. Indeed, *MYC* transcript abundance rose upon *CtBP1* silencing (Fig. 5C). Comparing the influence of TCF4 and CtBP1 on *MYC* expression, we found that *TCF7L2* silencing significantly down-regulated *MYC* (−0.9 log_2_FC), while *CtBP1* silencing significantly up-regulated *MYC* (0.73 log_2_FC). Given that *MYC* is a target gene of both TCF4 and LEF1, we investigated which WNT transcription factor is the primary regulator of *MYC*. Plotting the expression of *TCF7L2*, *LEF1*, and *MYC* over the siTCF7L2 or siLEF1 time series demonstrated that *MYC* expression more closely tracks the expression of *TCF7L2* than *LEF1* (Fig. 5D). Indeed, during the siTCF7L2 time series, while *TCF7L2* and *MYC* transcript abundance both fell, *LEF1* expression rose dramatically. The cooperation between TCF4 and CtBP1 to silence *LEF1*, the reliance of *MYC* expression upon TCF4, and the up-regulation of *MYC* upon *CtBP1* silencing, supports a competition centered on TCF4 transcriptional output.

## DISCUSSION

The canonical WNT signaling pathway maintains stemness in the intestines and is therefore necessary for the maintenance of the intestinal epithelium. Deletion of TCF4, the major WNT transcription factor in the intestines, compromises the intestinal stem cell compartment in neonatal and adult mice (*3*, *12*). Canonical WNT signaling activation manifests as an abundance of β-catenin, a transcriptional activator. Transmigration of β-catenin from the cytoplasm to the nucleus results in the formation of TCF/LEF and β-catenin complexes, which drive WNT target gene expression leading to cell cycle progression. Aberrant activation of the WNT signaling pathway is generally recognized to be the initiating event in colon cancer (*39–41*).

In the non-diseased human colon, *TCF7L1* and *TCF7L2* are the dominant WNT signaling transcription factors, while *TCF7* and *LEF1* are largely absent (*31*). This balance changes in colon cancer wherein *TCF7L1* is largely absent, while the relative expression of *TCF7* and *LEF1* increases with *TCF7L2* remaining the dominant WNT transcription factor (*42*, *43*). The high levels of nuclear β-catenin in colon cancer and the distinct roles played by the various WNT transcription factors suggests that a shift in relative WNT transcription factor abundance may alter the transcriptome, and therefore behavior, of colon cancer cells.

To investigate the impact of TCF4, the dominant WNT signaling transcription factor in the colon and colon cancer, we silenced *TCF7L2*, the gene encoding TCF4. *TCF7L2* silencing resulted in a robust, dose-dependent up-regulation of the WNT transcription factor *LEF1*, which translated to a similar increase in LEF1 nuclear protein abundance, supporting a WNT signaling intrinsic, self-regulatory mechanism. LEF1 was found to be transcriptionally competent, which may explain instances of increasing WNT activity and confirms previous observations of transcriptionally active LEF1 in colon cancer (*43*, *44*). The TCF4-*LEF1* interaction was observed in both a stable setting using CRISPR, as well as in multiple colon cancer cell lines including LoVo, SW480, DLD1, and RKO, but not HCT116, LS174T, or HT29. The differential *LEF1* response to *TCF7L2* silencing could be explained by the presence of the Catenin Inhibitory Domain (CID), a region of APC found to be critical for regulating β-catenin (*35*). However, CRISPR-induced mutation of the *APC* CID did not influence the TCF4-*LEF1* response in HCT116 or RKO, two lines originally harboring wild-type *APC*. Endogenous expression of the four WNT transcription factors in HCT116 and RKO, retrieved from the Cancer Cell Line Encyclopedia (CCLE), demonstrated that *LEF1* is not expressed in HCT116 but is the dominant WNT transcription factor in RKO, suggesting that an earlier process, perhaps during tumorigenesis, was responsible for establishment of the TCF4-*LEF1* interaction.

Since the APC::β-catenin::TLE axis appeared less effective at regulating *LEF1* expression than TCF4, we sought alternative WNT transcriptional regulators reported to interact with TCF4 and selected the CtBP family of transcriptional repressors (*45–49*). CtBP1 but not CtBP2 was found to repress *LEF1* expression in the β-catenin rich SW480 nucleus, as the silencing of *CtBP1*, but not *CtBP2*, resulted in *LEF1* up-regulation. However, LEF1 abundance only doubled upon CtBP1 reduction, while TCF4 reduction resulted in a 3-fold increase in LEF1 abundance. This indicates that *LEF1* was still partially repressed by TCF4. TCF4 also binds the TLE family of repressors, which may have supplemented TCF4-mediated repression. The use of multiple repressors suggests that TCF4 serves as a signal integrator from multiple pathways.

Since reduction of TCF4 or CtBP1 resulted in *LEF1* up-regulation, it appeared as if TCF4 and CtBP1 cooperate to repress *LEF1* expression. ChIP sequencing was performed for TCF4 and CtBP1 to determine whether *LEF1* regulation was direct or indirect. Clear overlap between TCF4 and CtBP1 DNA binding sites was observed at the *LEF1* promoter as well as along the *LEF1* gene body, supporting a model of direct *LEF1* regulation by TCF4 and CtBP1. However, it remains unclear whether TCF4 and CtBP1 bind one another or whether an additional binding partner is required for TCF4-CtBP1 repressive function. While TCF4 harbors domains for CtBP1 binding, experimental evidence is unclear (*48*, *50*).

Further analyses of CtBP1 ChIP-seq data revealed that CtBP1 exhibits DNA-binding patterns characteristic of a transcription factor, with considerable overlap (∼51%) between TCF4 and CtBP1 promoter-specific target genes. These findings suggest that TCF4 is a major relay for CtBP1-driven transcriptional repression. Additionally, a substantial fraction (∼40%) of CtBP1 binding occurred in promoters, verifying previous reports that CtBP1 is involved in short-range transcriptional repression (*51*). Short range repression is often mediated by histone deacetylases (HDACs), which have been reported to associate with CtBP1 (*52*). Notably, HDACs are known to interact with the TLE family of co-repressors, suggesting potential functional redundancy or cooperation between CtBP1 and TLE proteins in mediating WNT transcriptional repression.

The large overlap between CtBP1 and TCF4 regulatory sites prompted us to investigate the transcriptional impact of CtBP1 on SW480 cells in relation to the impact of TCF4. Silencing of *CtBP1* had a neutral impact on global gene expression patterns with a similar number of genes up- and down-regulated. *TCF7L2* silencing resulted in a net increase in global gene expression, suggesting an overall repressive role. GSEA demonstrated down-regulation of the MYC and E2F gene sets upon *TCF7L2* silencing, in line with the role of TCF4 in driving *MYC* expression and cell cycle progression. However, upon *CtBP1* silencing, MYC gene set activity increased, as did *MYC* expression, suggesting that CtBP1 represses *MYC*. CtBP1 ChIP-seq data identified a CtBP1-binding site within the *MYC* promoter, supporting a model of direct repression. Given that *MYC* expression decreases upon loss of TCF4, yet increases upon *CtBP1* silencing, suggests a competition focused on TCF4 transcriptional output. The balancing of TCF4 transcriptional activity may explain observations of TCF4 functioning as a tumor suppressor (*53–55*).

Since TCF4 or CtBP1 reduction increases the levels of transcriptionally competent LEF1 and given that *LEF1* is expressed in colon cancer but not non-diseased colon tissue, we silenced *LEF1* expression in SW480 to understand its impact on the colorectal cancer transcriptome. Broadly, more genes were significantly down-regulated than up-regulated, supporting the general role of LEF1 as an activator, contrasting from the overall pattern observed for TCF4. GSEA results for *TCF7L2* or *LEF1* silencing were largely non-overlapping indicating that these transcription factors drive different expression programs. *LEF1* silencing resulted in the down-regulation of immune associated gene sets such as TNFA Signaling, Inflammatory Response, and Interferon Alpha, following one of LEF1’s developmentally defined roles (*14*, *16*). Indeed, many gene sets down-regulated upon *LEF1* silencing, were up-regulated upon *TCF7L2* silencing, a situation wherein *LEF1* is up-regulated. Though despite the differences in GSEA, LEF1 and TCF4 cooperate to drive the expression of WNT target genes, such as *MYC*. However, TCF4 is the major regulator of *MYC*, not LEF1, since *TCF7L2* and *MYC* expression are strongly correlated and the up-regulation of *LEF1* upon *TCF7L2* silencing did not rescue *MYC* expression. This dependence of *MYC* upon TCF4 may underlie the dominance of TCF4 in colonic stem cells and colon cancer.

A limitation of our study is the reliance on established cell lines. Although cell lines provide a homogeneous and well-defined system well-suited for investigating mechanisms of transcriptional regulation, they do not recapitulate the *in vivo* cellular environment. As a result, transcriptional responses observed *in vitro* may differ from those occurring *in vivo*. A second limitation of our study is the CtBP1 ChIP-seq data. Given that CtBP1 accesses the DNA indirectly through binding partners, we sought to capture those DNA regions at which CtBP1 is localized by crosslinking DNA-protein and protein-protein interactions. As a result of the additional crosslinking interactions necessary to capture CtBP1 binding events, the CtBP1 ChIP-seq data suffers from increased background noise and, while useful, should be interpreted with caution.

In summary, we find that TCF4 cooperates with CtBP1, a transcriptional repressor, to directly repress *LEF1* expression in colon cancer. Reduction of TCF4 or CtBP1 increases the amount of transcriptionally competent LEF1, which drives a different transcriptional program than TCF4, favoring immune associated gene sets (Figure 6). While TCF4 and LEF1 both activate *MYC* expression, TCF4 is the dominant *MYC* activator. However, the interaction of TCF4 with CtBP1 moderates *MYC* expression, which is elevated upon *CtBP1* silencing. These findings characterize the role of LEF1 in colon cancer, support a dual activator-repressor role for TCF4 in colon cancer, and identify a CtBP1-dependent avenue for mitigating *MYC* expression.

**Figure 6.**
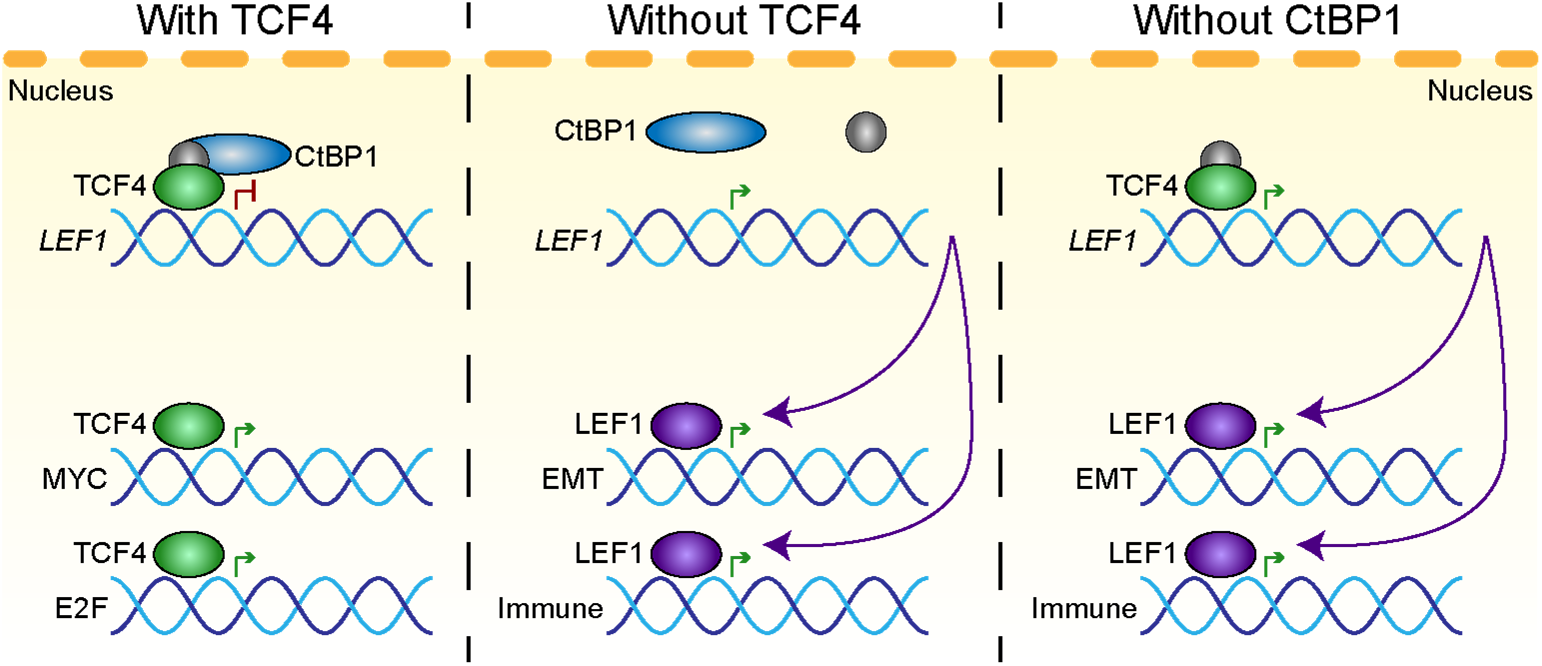
TCF4, CtBP1, and LEF1 in Colon Cancer|. Diagram showing the proposed interaction between TCF4, CtBP1, and *LEF1*. When TCF4 is present it forms a complex with CtBP1 (left), possibly with the assistance of an additional unidentified protein (grey). The TCF4 and CtBP1 complex binds the *LEF1* promoter and represses expression. TCF4 also drives the expression of genes within the MYC and E2F gene sets. Without TCF4 (middle), CtBP1 does not bind to the *LEF1* promoter, and *LEF1* expression is increased three-fold. Increased LEF1 leads to the up-regulation of genes in the EMT and Immune-associated gene sets. Without CtBP1 (right), TCF4 can still bind the *LEF1* promoter but repression is only partially effective and *LEF1* expression increases two-fold.

## MATERIALS AND METHODS

### Cell Culture

The DLD1, HCT116, HT29, LoVo, LS174T, RKO, and SW480 colon cancer cell lines were purchased from the American Type Culture Collection (ATCC). DLD1 and SW480 were cultured in RPMI-1640 (Thermo Fisher; 11875093). HCT116 and HT29 were cultured in McCoy’s 5A Modified Medium (Thermo Fisher; 16600082). LoVo was cultured in Ham’s F-12 (Thermo Fisher; 11765054), LS174T was cultured in Minimum Essential Medium (Thermo Fisher; 11095080), while RKO was cultured in Dulbecco’s Modified Eagle Medium (Thermo Fisher, 11965092). All growth media were supplemented to 10% FBS (Thermo Fisher; 16140071) and 2 mM L-Glutamine (Thermo Fisher; 25030081). Cell lines were grown in T75 Culture Flasks (Corning; 430641U) at 37°C in a humidified environment containing 5% CO_2_. Cell line identity, assessed via STR Profiling, and the absence of mycoplasma contamination, assessed via PCR (Genlantis; MY01050), was confirmed every six months.

The treatment-naïve rectal cancer biopsy, single cell-derived cell line (SCDCL) was grown in F-medium consisting of 25% Ham’s F-12, 75% DMEM, supplemented to 10% FBS, 2 mM L-Glutamine, 100 μg/mL Streptomycin and 100 Units/mL Penicillin (Thermo Fisher; 15140122), 25 ng/mL Hydrocortisone (Millipore Sigma; H0888), 5 mg/mL Insulin (Millipore Sigma; I5500), 0.1 nmol/L Cholera Toxin (Millipore Sigma; C3012), 250 ng/mL Amphotericin B (Fisher Scientific; BP264550), 0.125 ng/mL Human Epidermal Growth Factor (EGF) Recombinant Protein (Thermo Fisher; PHG0313), 10 μg/mL Gentamicin (Thermo Fisher; 15710064), and 5 mmol/L ROCK inhibitor Y-27632 (Enzo; 270333M025). Cells were cultured at 37°C in a humidified incubator with 5% CO_2_ and 2% oxygen.

### Inverted siRNA Transfections

SW480 cells were seeded in a 6-well tissue culture plate (Corning; 353046) at a density of 150,000 cells per well. For cell types that grew more quickly, such as DLD1, the starting cell density was 50,000 cells per well. The wells contained 2 mL of the appropriate growth medium fortified to 10% FBS and 2 mM L-Glutamine. Silencer Select siRNAs (Thermo Fisher) Negative Control #1 (4390843), siTCF7L2-s13880, siLEF1-s27618, siCtBP1-s3699, or siCtBP2-s3702 were delivered to the cells using the Lipofectamine RNAiMAX Transfection Reagent (Thermo Fisher; 13778150) and Opti-MEM Medium (Thermo Fisher; 31985070) according to the manufacturer’s instructions. Additional details regarding the transfection methodology are available (*56*).

### CRISPR/Cas9

Single guide RNAs (sgRNAs) targeting *TCF7L2* were designed using CRISPick which flanked the exons encoding the HMG-box domain. The sgRNAs were cloned into pSpCas9(BB)-2A-Puro (PX459) V2.0 (Addgene; #62988) as previously described (*57*) and successful sgRNA integration was confirmed with Sanger sequencing. pSpCas9(BB)-2A-Puro-*TCF7L2* plasmids were delivered to SCDCL using Lipofectamine 3000 (Thermo Fisher; L3000008), according to the manufacturer’s instructions. Puromycin (InvivoGen; ant-pr-1) selection for transfected cells began 48 hours after transfection and continued for seven days. After assessing the CRISPR efficiency by PCR, the transfected population was single-cell sorted and expanded for at least 2 weeks. Following expansion, the *TCF7L2*-CRISPR single cell clones were subjected to PCR genotyping to confirm *TCF7L2* knockout. The sgRNA and genotyping primer sequences can be found in Supplementary Table 1.

### Quantitative PCR

RNA was extracted from at least three independent, inverted transfections using the RNeasy Mini Kit and on-column DNase treatment (Qiagen; 74104, 79254). RNA concentration was determined using a NanoDrop 1000 spectrophotometer (Thermo Fisher). Synthesis of cDNA was performed with 800 ng of RNA using the Verso cDNA Synthesis Kit (Thermo Fisher; AB1453B). The optional RT Enhancer was used and equal amounts of both Anchored Oligo dT and Random Hexamers were added. The cycling protocol consisted of a 30-minute incubation at 42°C, a 2-minute incubation at 95°C, and was concluded with a 4°C hold. Quantitative PCR was performed for *TCF7L2*, *LEF1*, *AXIN2*, and *YWHAZ* on an ABI PRISM 7000 Sequence Detection System (Applied Biosystems) or a Lightcycler 480 II (Roche) using *Power* SYBR Green PCR Master Mix (Thermo Fisher; 4367659). The thermocycling protocol consisted of a 2-minute incubation at 50°C, which proceeded into a 10-minute incubation at 95°C, followed by 40 two-step cycles consisting of 15 seconds at 95°C and 1 minute at 60°C. The cycling protocol was concluded with a melting curve analysis. Fold change was determined using *YWHAZ* as the reference gene and the 2^-ΔΔCT^ method (*58*). Primer sequences are provided in Supplementary Table 2.

### Western Blot

Nuclear and cytoplasmic protein fractions were extracted from SW480 cells using the NE-PER Nuclear and Cytoplasmic Extraction Reagents (Thermo Fisher; 78833). A brief sonication step consisting of five, 1 second ON: 2 seconds OFF cycles performed with a Branson 450 Digital Sonifier (Branson Ultrasonics) at 10% Duty Cycle was used to disrupt nuclei following digestion with NER. Protein was prepared for quantification using the Pierce BCA Protein Assay Kit (Thermo Fisher; 23225) and quantified on a SpectraMax M2^e^ microplate reader (Molecular Devices). Western blot samples were generated with 30 µg protein, NuPAGE LDS Sample Buffer (Thermo Fisher; NP0007), and Molecular Biology Grade Water (Corning; 46000CM). Samples were heated to 70°C for 10 minutes. The samples, 25 µL in volume, were electrophoresed for 50 minutes at 200V through a 4-12% Bis-Tris gel (Thermo Fisher; NP0321BOX) in an XCell *SureLock* Mini-Cell (Thermo Fisher; EI0001) with MOPS Running Buffer (Thermo Fisher; NP0001). Proteins were transferred into an Immobilon-P PVDF Membrane (Millipore; IPVH00010) for 90 minutes at 30V using Transfer Buffer (Thermo Fisher; NP00061) and an XCell II Blot Module (Thermo Fisher; EI9051). Membranes were blocked in 1X TBS (Takara, T903) with 5% milk (VWR; M20310G) for 1 hour at room temperature, shaking. Primary antibody incubations were performed overnight, shaking, at 4°C in 1X TBS with 5% BSA (Roche; 03117332001) using dilutions recommended by the manufacturer (usually 1:1000). Primary antibodies used in this study targeted TCF4 (CST; 2569S), LEF1 (CST; 2230S), CtBP1 (CST; 8684S), CtBP2 (CST; 13256S), or TBP (CST; 8515S). HRP-linked secondary antibody incubations were performed in 1X TBS for 1 hour at room temperature, using dilutions recommended by the manufacturer (1:2000). The secondary antibody used in this study was Goat Anti-rabbit IgG, HRP-linked (CST; 7074S). Detection was performed using SuperSignal West Pico PLUS Chemiluminescent Substrate (Thermo Fisher; 34580) for 5 minutes in the dark. Blots were imaged using an Azure c600 Gel Imaging Station (Azure Biosystems) and exposure time was optimized for capture of faint bands.

### Dual-Luciferase Assay

SW480 cells were seeded in an opaque, white 96-well tissue culture plate (Corning; 3917) at a density of 5,000 cells per well. Cells were transfected 24 hours later with the Cignal TCF/LEF Reporter Assay Kit (Qiagen; CCS018L), using Lipofectamine 3000 (Thermo Fisher; L3000008). Cells were simultaneously transfected with siNeg, siTCF7L2, or siTCF7L2 + siLEF1, according to the Inverted Transfection protocol, using RNAiMAX. Cell lysates were prepared after the time course had been completed using the Dual-Luciferase Reporter Assay System (Promega; E1910), based on the manufacturer’s instructions for Passive Lysis. Twice the recommended amount of 1X PLB was added per well. Luciferase luminescence was measured according to the Dual-Luciferase Reporter Assay System instructions, using a Centro XS^3^ LB 960 Microplate Luminometer (Berthold Technologies). Specifically, 100 μL of LAR II was injected into a well, followed by a 2-second delay, followed by a 10-second measurement. Immediately thereafter, 100 μL of Stop & Glo reagent was injected, followed by a 2-second delay, and concluded with a final 10-second measurement. Each biological replicate consisted of three wells containing TCF/LEF Reporter Plasmid, one well containing Negative Control, one well containing Positive Control, and one well containing Non-Transfected Control for each sample. Data was analyzed by determining the Firefly to Renilla Luciferase ratio, then calculating the fold change of the F:R ratios against Time 0.

### Immunocytochemistry

Parental or Homozygous *TCF7L2* knockout SCDCL cells were grown on coverslips, rinsed with 1X PBS (Thermo Fisher; 10010023) for 5 minutes, and fixed with 4% paraformaldehyde (Fisher Scientific; AAJ19943K2) for 10 minutes at room temperature. Cells were then washed with 1X PBS and incubated with 0.5% Triton X-100 (Millipore Sigma; X100) in 1X PBS for 10 minutes at room temperature. Cells were then washed with 1X PBS and blocked in 5% BSA for 1 hour at room temperature. Cells were incubated overnight at 4°C in 5% BSA with the primary antibody diluted according to the manufacturer’s recommendations (1:200). The following morning, cells were washed with 1X PBS and incubated for 2 hours at room temperature in the dark with secondary antibody diluted according to the manufacturer’s recommendations (1:1000). The secondary antibodies used were Goat anti-Rabbit Alexa Fluor Plus 488 (Thermo Fisher; A32731TR) or Goat anti-Mouse Alexa Fluor Plus 555 (Thermo Fisher; A32727). After the final PBS wash, coverslips were mounted with ProLong Gold Antifade with DAPI (Thermo Fisher; P36931).

### RNA Sequencing

Three independent inverted transfections were performed in SW480 followed by RNA extraction using the RNeasy Mini Kit and on-column DNase treatment. RNA concentration and integrity was determined using a TapeStation 4200 (Agilent). RNA samples with an RNA Integrity Number (RIN) of 9 or higher were sent to the NCI CCR Sequencing Facility for library preparation and sequencing. Libraries were prepared using the TruSeq Stranded RNA Library Prep Kit (Illumina). Libraries were sequenced on either a NextSeq 550 or a Novaseq 6000 generating ∼50 million, 150 base pair, paired-end reads per library. Reads were demultiplexed using Bcl2fastq (v2.20) and adaptor/quality trimming was performed using Cutadapt (v1.18). Alignment to hg38 was performed using STAR (v2.7.0f). General RNA statistics were calculated using Picard (v2.18.26) and quantification was performed using RSEM (v1.3.1). Differential gene expression analysis was performed using the appropriate CCBR pipeline. Many figures were generated using the R tidyverse (*59*), while volcano plots were generated using the EnhancedVolcano package (*60*). Gene Set Enrichment Analysis (GSEA) was performed using the fgsea package and the Hallmark gene sets from MSigDB (*61*).

### ChIP Sequencing

Chromatin Immunoprecipitation was performed in SW480 with the ChIP-It High Sensitivity Kit (Active Motif; 53040). Chromatin was sheared by sonication with a Branson 450 Digital Sonifier (Branson Ultrasonics) for a total of 3 minutes and 30 seconds ON time, split into 30 second ON – 30 second OFF cycles at 15% amplitude. For each immunoprecipitation, 20 µg of sheared chromatin was mixed with 1 µg of antibodies targeting TCF4 (CST; 2569S) or 1 µg of antibodies targeting CtBP1 (CST; 8684S). Steps not described here were performed according to the manufacturer’s instructions. Immunoprecipitated DNA quality and size was determined using a TapeStation 4200 (Agilent). Samples were sent to the NCI CCR Sequencing Facility for library preparation and sequencing. Libraries were prepared using the Swift 2S DNA Library Prep Kit (Swift Biosciences). Libraries were sequenced on a Novaseq 6000 generating ∼50 million, 100 base pair, single-end reads per sample. Reads were demultiplexed using Bcl2fastq (v2.20) and adaptor/quality trimming was performed using Cutadapt (v1.18). Alignment to hg38 was performed using Bowtie2 (v2.3.4.1) and general mapping statistics were performed using Picard (v2.18.26). Peak calling and motif enrichment analysis was performed using the appropriate CCBR pipeline. Peak heatmaps were generated using deepTools (*62*).

### Statistical Analysis

To compare between different groups, a two-tailed Student’s *t*-test was used, if not otherwise stated, with * denoting *p* < 0.05 and ** denoting *p* < 0.01. In instances where *p* < 0.001, ** was also used. Comparisons involving multiple timepoints were compared to the Time 0 measurement, while comparisons with multiple cell lines used the Time 0 values from the respective cell line. Calculations with the truncated APC lines were compared to the Time 0 measurement from the wild-type cell line. Data analysis of RNA-seq and ChIP-seq data was performed in R.

### Data Retrieval and Filtering – GTEx and TCGA

Data from the Genotype-Tissue Expression project (GTEx) was accessed using *recount3* (*63*). Samples collected must have been scored as a 2 or lower on the Hardy Scale, the RNA Integrity Number (RIN) must have been 7 or higher, the proportion of exonic reads among uniquely mapping reads must have been 70% or higher, all samples must have been sequenced using TruSeq.v1 chemistry, and all samples listed in sample tissue changes were removed.

Data from The Cancer Genome Atlas (TCGA) was accessed using *recount3*. Samples must have originated from the primary tumor, the histological type must have been Colon Adenocarcinoma, the percent of tumor cells must have been 70% or greater, the percent of normal cells must have been 20% or less, and the percent of stromal cells must have been 20% or less. Tumor purity data were retrieved from (*64*).

## Acknowledgments

We thank Dr. Thomas Ried and Dr. Kan Cao for their insight and advice throughout the project. We also thank the members of the Collaborative Bioinformatics Resource of the Center for Cancer Research of the NCI for their guidance and support. This work utilized the computational resources of the NIH HPC Biowulf cluster. This research was supported in part by the Intramural Research Program of the National Institutes of Health (NIH).

The contributions of the NIH author(s) are considered Works of the United States Government. The findings and conclusions presented in this paper are those of the author(s) and do not necessarily reflect the views of the NIH or the U.S. Department of Health and Human Services.

## Author contributions

Conceptualization: MAB, TZ, MB, JC, PM

Methodology: MAB, SY, JK, WDC

Investigation: MAB, SY, JK, WDC, DW, STK

Visualization: MAB, SY

Supervision: TZ, MB, JC, PM

Writing - original draft: MAB

Writing - review & editing: SY, TZ, MB, JC, PM

## Data Accessibility Statement

Data has been submitted to the Gene Expression Omnibus (GEO) under the accession numbers: GSE331304, GSE331307, and GSE331308.

## Supplementary Figures and Materials

**Supplementary Figure 1.**
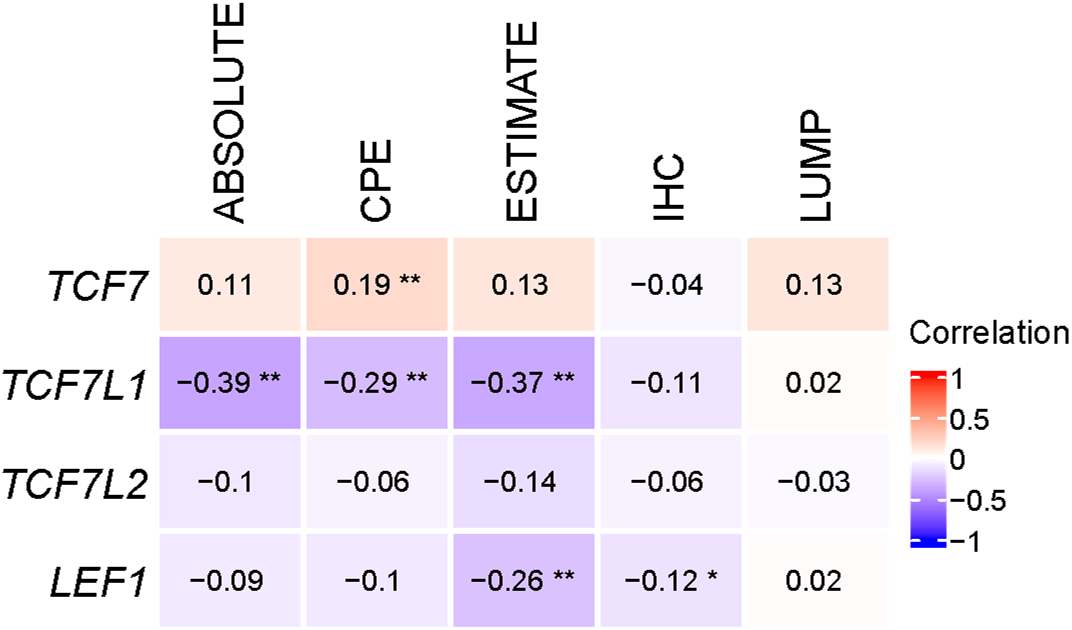
Tumor Purity and WNT Transcription Factor Correlations |. Tumor purity metrics were retrieved and correlated to the expression (TPM) of the WNT transcription factors (*64*). The Pearson correlation coefficient is denoted by the color and displayed in the box (Student’s *t*-test, * *p* < 0.05, ** *p* < 0.01).

**Supplementary Figure 2.**
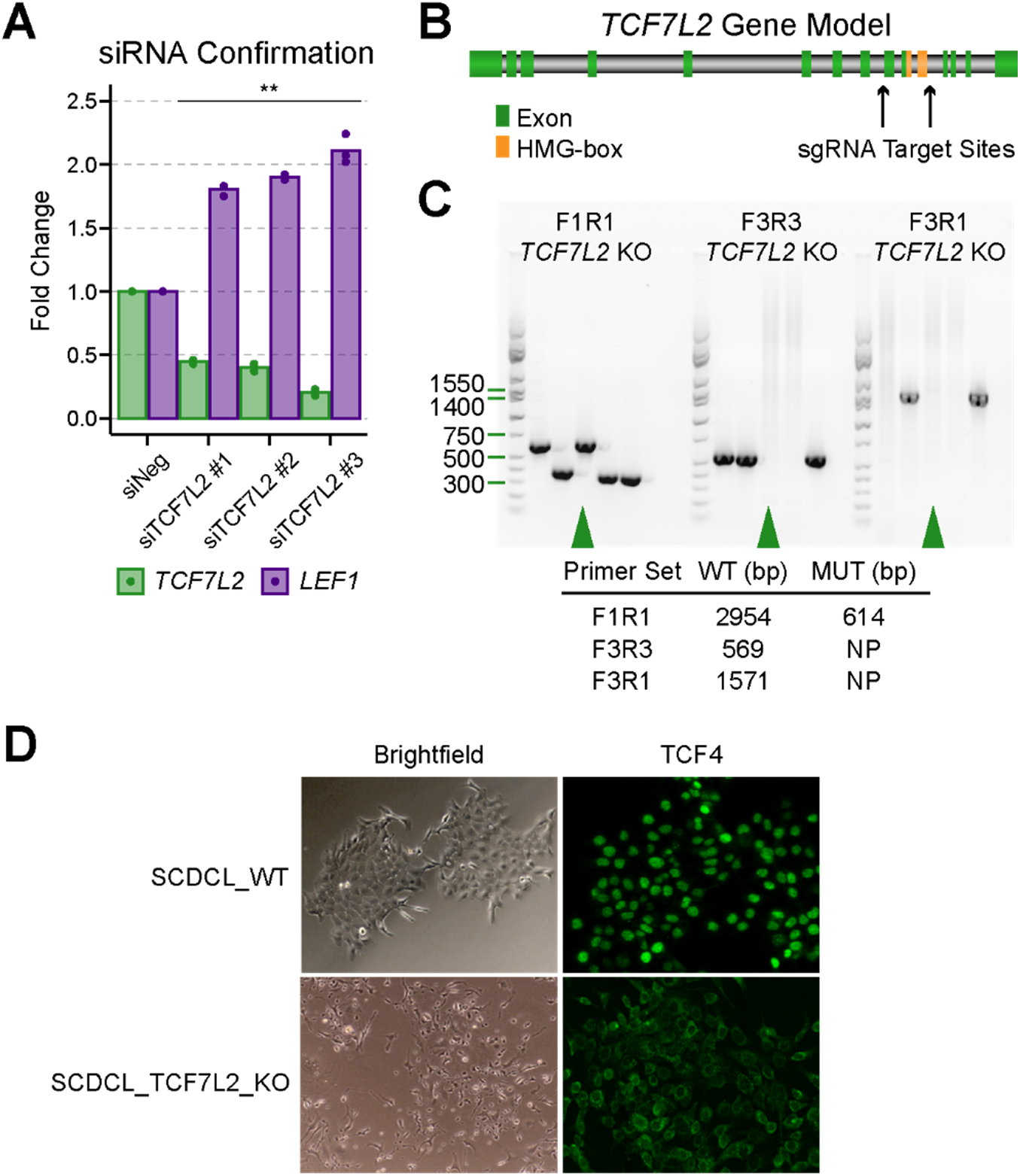
siTCF7L2 and Dose-Dependent *LEF1* Up-regulation. **| A |** Different siRNAs targeting various regions of *TCF7L2* were assessed for silencing efficiency and *LEF1* up-regulation as measured by qPCR at the 72-hour timepoint. The siRNAs silenced *TCF7L2* with varying efficiency, which resulted in a dose-dependent up-regulation of *LEF1* (Student’s *t*-test, ** *p* < 0.01). Three biological replicates are plotted. **| B |** Gene model of *TCF7L2* indicating the relative position of exons, the DNA binding HMG-box, and the sgRNA target sites. **| C |** PCR-based genotyping of *TCF7L2* knockout (KO) lines to assess successful removal of the HMG-box domain. **| D |** Immunofluorescence of the Parental *TCF7L2* WT SCDCL and Homozygous *TCF7L2* KO SCDCL lines. Note the diffuse TCF4 (*TCF7L2)* staining in the *TCF7L2* KO.

**Supplementary Figure 3.**
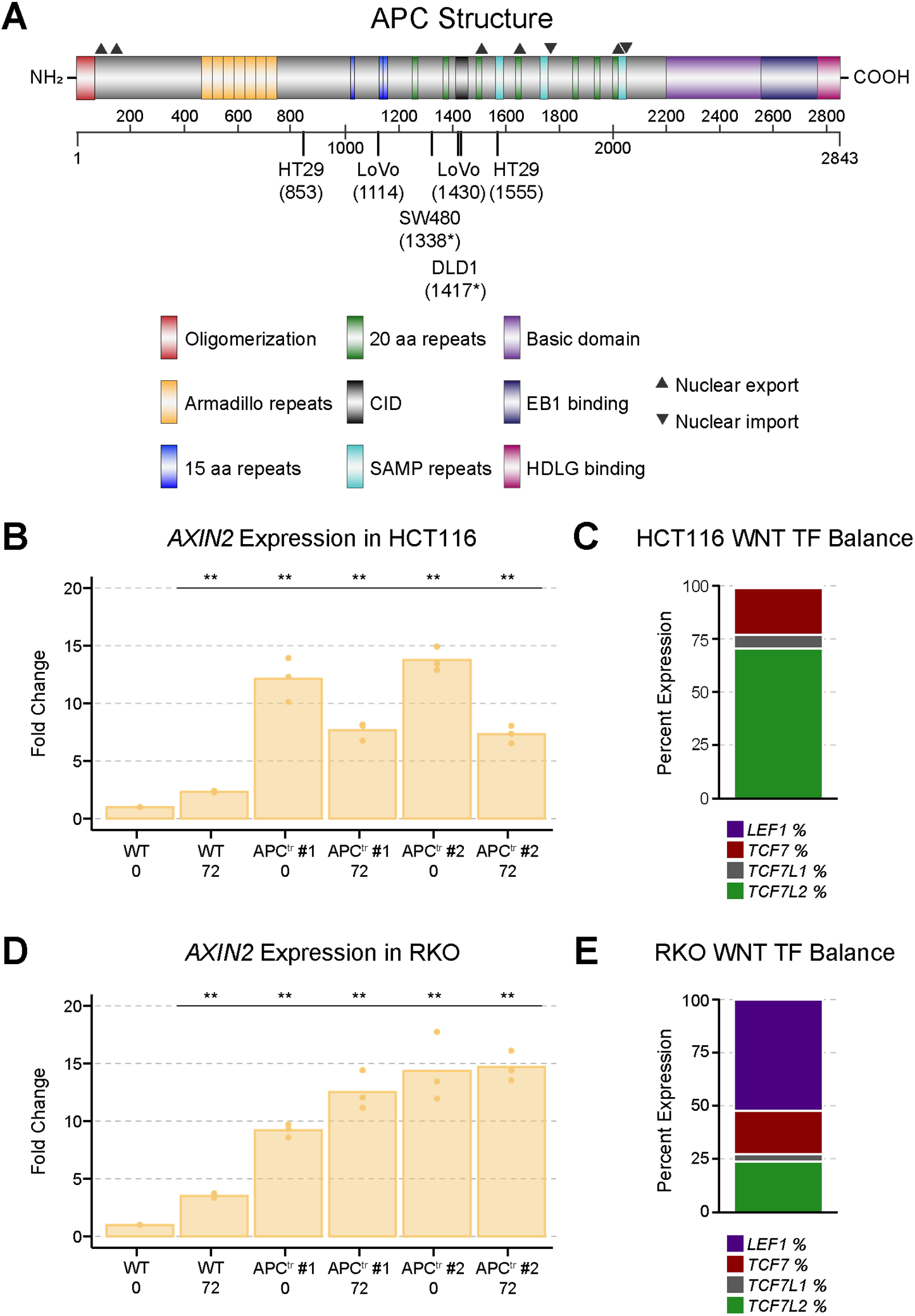
*AXIN2* Expression is Deregulated Upon APC Truncation. **| A |** An APC protein model with functional domains provided as well as amino acid numbers. Colon cancer cell lines used in this study are placed at the position of the *APC* mutations they harbor, specified in parentheses. An asterisk (*) next to the amino acid number indicates a loss of heterozygosity. **| B |** *AXIN2* expression in wild-type HCT116 and APC truncated clones upon silencing of *TCF7L2* at the 72-hour timepoint. Fold change was calculated using the siNeg control (WT 0). Three biological replicates are plotted (Student’s *t*-test, ** *p* < 0.01). **| C |** HCT116 RNA sequencing data from the Cancer Cell Line Encyclopedia (CCLE), normalized to TPM, was used to generate the WNT transcription factor expression percentage, which is the percent that each WNT transcription factor contributes to the total WNT transcription factor expression. *LEF1* is not expressed in HCT116. **| D |** *AXIN2* expression in wild-type RKO and APC truncated clones upon silencing of *TCF7L2* at the 72-hour timepoint. Fold change was calculated using the siNeg control (WT 0). Three biological replicates are plotted (Student’s *t*-test, ** *p* < 0.01). **| E |** RKO RNA sequencing data from the CCLE, normalized to TPM, was used to calculate the WNT transcription factor balance. *LEF1* is the dominantly expressed WNT transcription factor in RKO. Upon *TCF7L2* silencing in RKO, *LEF1* expression increases further.

**Supplementary Figure 4.**
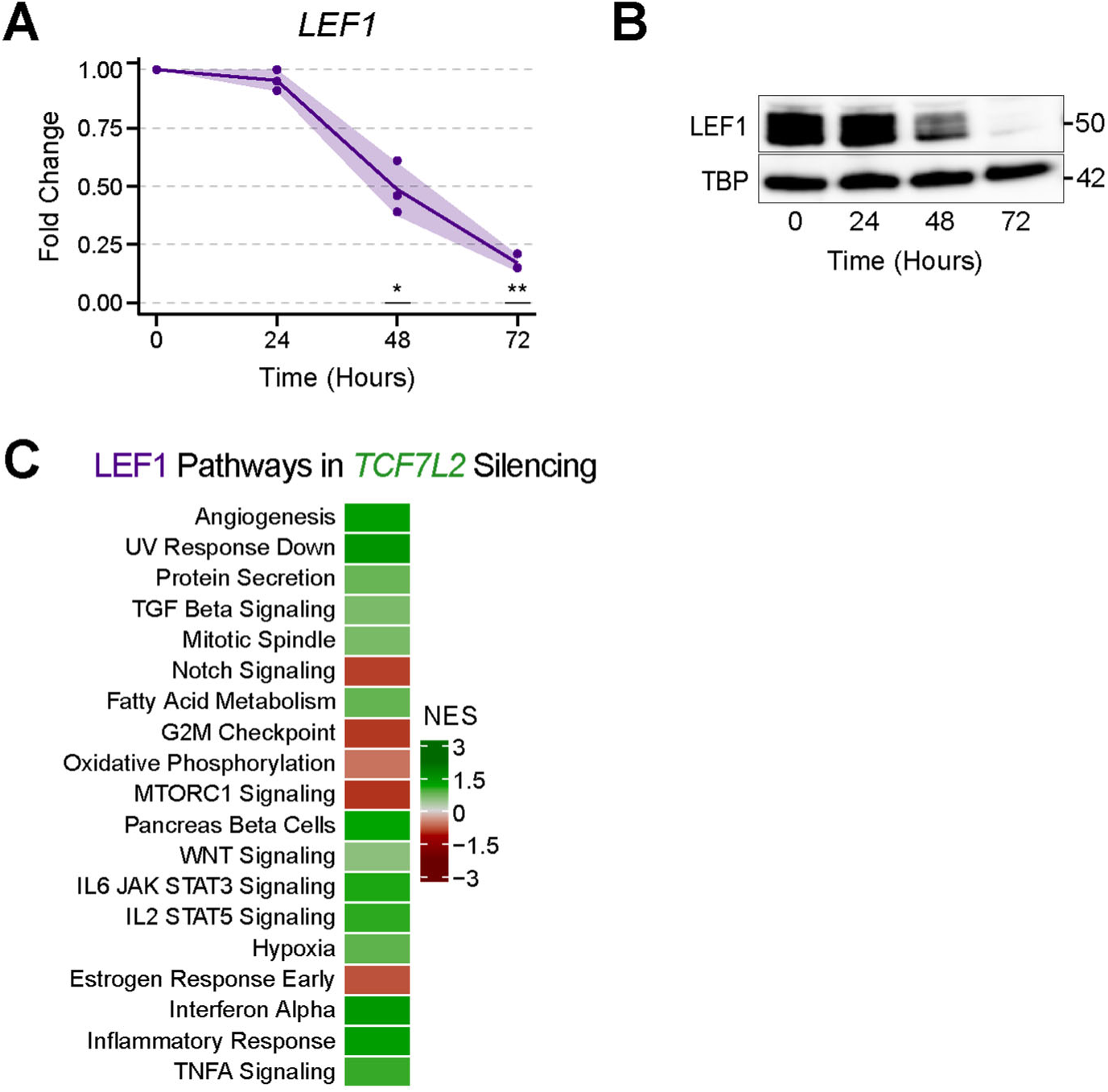
*LEF1* Expression Positively Correlates with Immune System Gene Set Activity. **| A |** Quantitative PCR (qPCR) was used to assess *LEF1* transcript abundance upon transfection with siLEF1 over the time series in SW480. Three biological replicates are plotted, the line represents the average, and the ribbon denotes the standard deviation (Student’s *t*-test, * *p* < 0.05, ** *p* < 0.01). **| B |** Nuclear LEF1 abundance during the siLEF1 time series was determined using Western Blot with TATA-Binding Protein (TBP) as loading control. LEF1 abundance decreased by ∼98% by the 72-hour timepoint. **| C |** The deregulated pathways upon *LEF1* silencing were plotted using the siTCF7L2 data to determine whether the down-regulated pathways upon *LEF1* silencing are up-regulated upon *TCF7L2* silencing (which induces *LEF1* over-expression). Indeed, 8 out of 10 down-regulated pathways upon *LEF1* silencing are up-regulated when *LEF1* is over-expressed (*TCF7L2* silencing).

**Supplementary Table 1.**
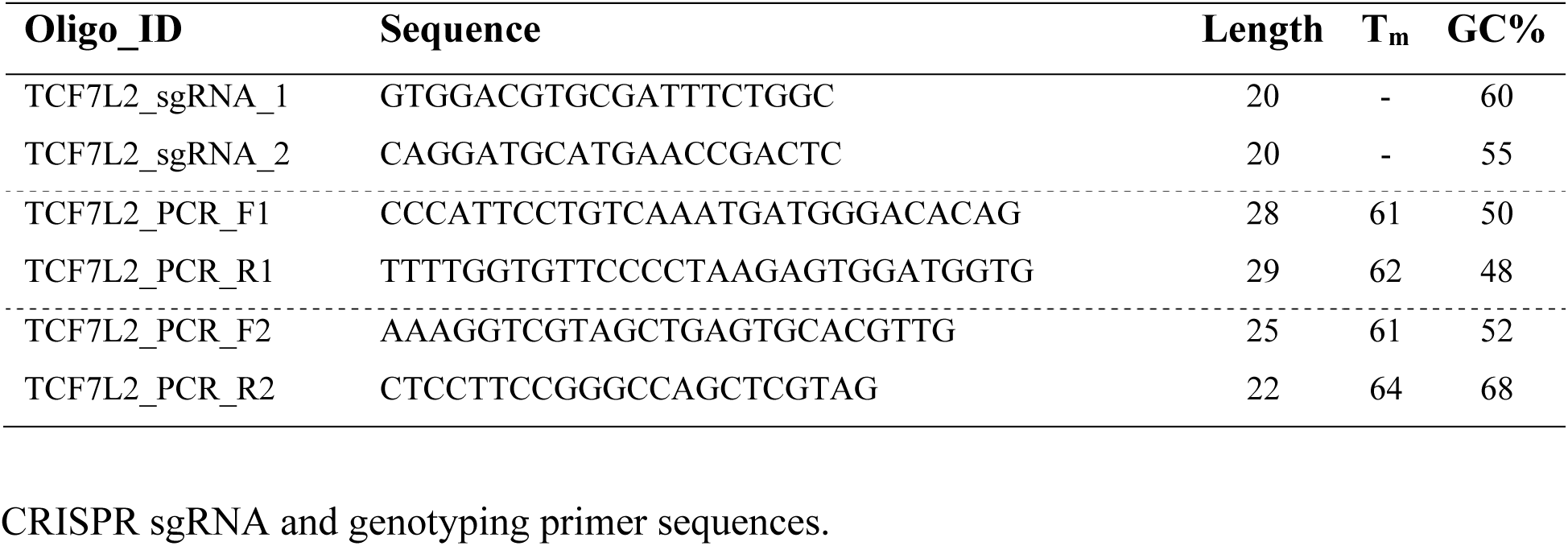
CRISPR sgRNA and genotyping primer sequences.

**Supplementary Table 2.**
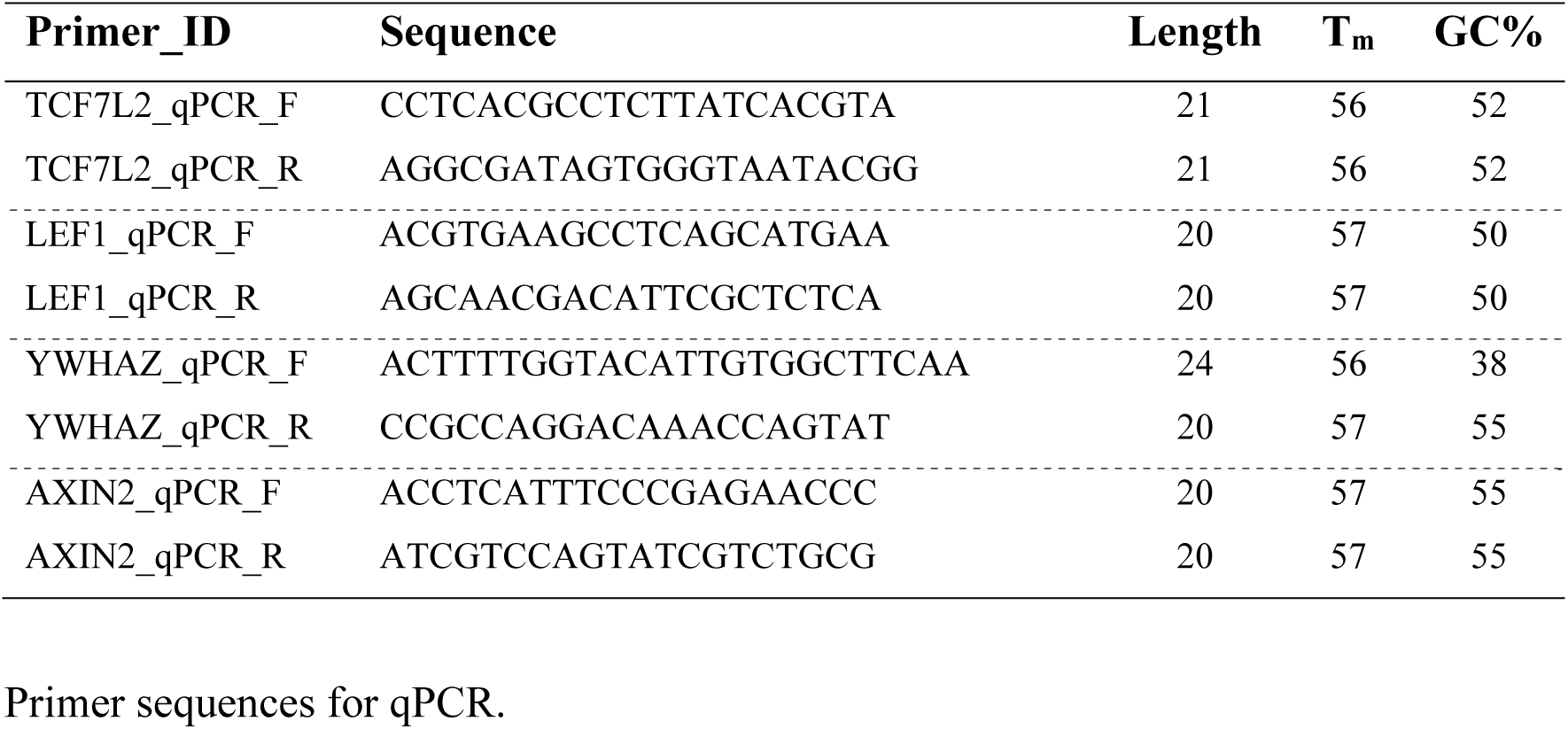
Primer sequences for qPCR.

